# Spared corticospinal neurons activate an endogenous plasticity program after partial CNS injury

**DOI:** 10.64898/2026.09.08.750239

**Authors:** Matias Murillo, Ciara F O’Brien, Noa Golan, Emma X Yin, Jessica Mariani, Michael P J Deans, Novin Balafkan, Kristen J Brennand, William B J Cafferty

## Abstract

Functional recovery after incomplete spinal cord injury depends substantially on the capacity of anatomically spared neurons to remodel their connections, yet the molecular programs underlying this endogenous plasticity remain poorly understood. To identify and validate these mechanisms, we combined retrograde labeling, unilateral corticospinal tract injury, and spatial transcriptomics in mice, with an *in vitro* human neuron model of neurite regeneration. Injured corticospinal neurons (CSNs) exhibited widespread transcriptional downregulation; the few activated pathways included stress, cell death, and degenerative programs. In contrast, spared CSNs activated a coordinated pro-plasticity program characterized by metabolic, immune, and cytoskeletal remodeling together with suppression of axon growth-inhibitory signaling. Computational network and drug perturbation analyses identified, and human neuronal experiments validated, *ARHGEF12* suppression and the histone deacetylase inhibitor (HDAC) vorinostat as complementary target- and state-based strategies to enhance neurite regeneration. These findings define key components of endogenous plasticity and provide a framework for discovering neural repair targets.

## Introduction

Stroke, traumatic brain injury, and spinal cord injury (SCI) are leading causes of long-term disability, reflecting the limited capacity of the adult central nervous system (CNS) for axon regeneration^1^. Nevertheless, a modest degree of spontaneous functional recovery can occur following incomplete injuries, largely attributed to remodeling of anatomically spared circuits^2–4^. Previous transcriptomic studies using single-nucleus RNA sequencing (snRNA-seq), translating ribosome affinity purification sequencing (TRAP-seq), and Patch-seq identified transcriptional programs associated with axotomy, regeneration failure, and regenerative interventions^5–7^. These studies have provided important insight into the molecular response of injured corticospinal neurons (CSNs) following CNS injury. In contrast, comparatively little is known about the molecular programs that enable spared neurons to undergo adaptive plasticity and establish new functional connections. Defining these endogenous repair mechanisms could reveal therapeutic strategies that enhance recovery without requiring long-distance regeneration of injured axons.

We demonstrated that actively sprouting CSNs exhibit a distinct transcriptional program that can be leveraged to enhance corticospinal remodeling and functional recovery after injury^4^, but this analysis was limited to a small population of CSNs profiled at a single late post-injury time point in *Ngr1* null mice. Consequently, it remains unknown whether anatomically spared CSNs more broadly engage a coordinated endogenous molecular repair response after injury in wildtype animals. Moreover, whereas we and others previously profiled intact CSNs post-injury using fluorescence-activated cell sorting and single-cell RNA sequencing^8–11^, recent advances in spatial transcriptomics now permit high-resolution and genome-wide interrogation of gene expression^12,13^ of sprouting CSNs within intact tissue and without dissociation-induced transcriptional artifacts. This approach provides an opportunity to directly compare the transcriptional states of injured and spared CSNs.

Functional validation and clinical translation of candidate therapeutic targets remain important challenges in CNS regeneration research. Despite substantial advances in preclinical studies with animal models, relatively few restorative therapies have successfully been translated into clinical practice^14–16^. Important differences in corticospinal tract (CST) anatomy, circuit organization, and neuronal development between humans and rodents raise the possibility that mechanisms governing axon growth and repair are not fully conserved across species^17–19^. Human induced pluripotent stem cell (hiPSC)-derived cortical neurons provide a scalable and genetically tractable platform for modeling neurite regeneration in a human genetic background. Integrating transcriptomic discovery *in vivo* with functional validation in hiPSC-derived neurons therefore established a complementary framework for identifying and prioritizing therapeutic strategies with greater translational potential.

Here, we combined retrograde labeling, unilateral CST injury and spatial transcriptomics to define the transcriptional programs of injured and spared CSNs. We identified a coordinated endogenous pro-plasticity signature in spared neurons that was distinct from the injury response and leveraged this signature to prioritize two independent interventions. First, genetic suppression of *ARHGEF12*, a component of the RhoA growth-inhibitory signaling cascade that was consistently among the most downregulated genes in spared neurons. Second, a pharmacological treatment, the HDAC-inhibitor vorinostat, predicted to induce a gene expression profile that mimicked the endogenous plasticity program. Both genetic and pharmacological strategies successfully increased neurite regeneration in hiPSC-derived cortical neurons. Together, these findings define key components of endogenous plasticity in response to CNS injury and establish an integrative framework for discovering and validating therapeutic targets for neural repair.

## Results

### Integrated spatial transcriptomic strategy enables comparative profiling of injured and spared CSNs

To identify and compare the transcriptional states of injured and spared corticospinal neurons (CSNs), we designed an experimental strategy combining retrograde labeling, unilateral corticospinal tract injury (pyramidotomy uPyX) and spatial transcriptomics (**Figure 1**). CSNs were labeled via bilateral injections of AAVrg-CAG-tdTomato into the cervical or lumbar spinal cord two weeks prior to uPyX (**Figures 1A, 1B**). The uPyX selectively transects corticospinal tract (CST) axons at the level of the medullary pyramids, without damaging the cell bodies or local cortical circuitry^4^. Crucially, the unilateral nature of the injury specifically targets CSNs from the ipsilateral cortex while sparing the contralateral side (**Figure 1C**). Visium spatial RNA-sequencing was performed on coronal sections of primary motor cortex (M1) 10 days after injury (**Figure 1A**). Injured and spared CSNs within the same cortex enabled direct within-animal comparisons, while the sham animal sections served as baseline for intact gene expression and between-animal comparison. Histological assessment confirmed robust bilateral labeling of CSN populations and preserved tissue morphology (**Figure 1D**). This strategy enabled characterization of gene expression changes in injured and spared CSNs, within their native cortical environment.

**Figure 1.**
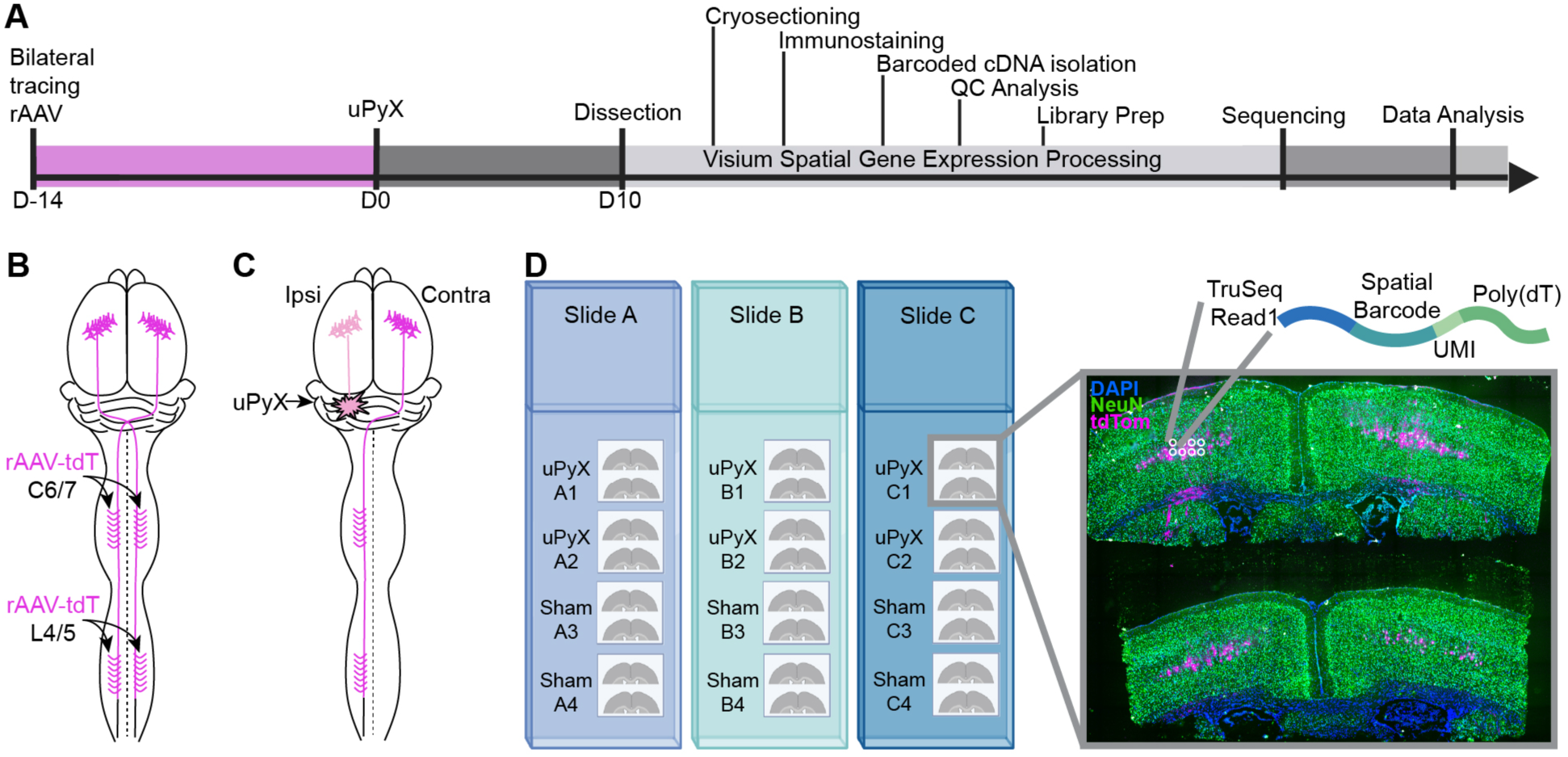
Integrated spatial transcriptomic strategy enable comparative profiling of injured and spared CSNs. **(A)** Experimental timeline. **(B)** Bilateral retrograde labeling for cervical and lumbar CSNs. **(C)** Injury model. Unilateral pyramidotomy (uPyX) selectively transects corticospinal tract axons originating from the ipsilateral hemisphere while preserving contralateral CSNs. **(D)** Distribution of uPyX and sham tissue sections across three Visium slides (Slides A–C). Inset: Representative M1 sections showing DAPI (blue), NeuN (green), and tdTomato-labeled CSNs (magenta) positioned over the Visium capture area. Inset illustrates Visium spots containing oligonucleotides with sequencing adaptor, spatial barcode, unique molecular identifier (UMI), and poly(dT) region.

Next, we assessed sequencing quality, library complexity, and reproducibility, confirming low mitochondrial and hemoglobin reads and robust transcript capture (**Figure S1**). This quality control and clustering pipeline demonstrated consistent sequencing performance and anatomically resolved gene expression patterns across biological replicates. To identify cell-type composition in M1 and isolate CSN-enriched spots for downstream analysis, we performed deconvolution aided by retrograde labeling and integration of single-cell RNA sequencing (scRNAseq) data (**Figure 2**). On average the 55µm spots contained 3-7 nuclei (**Figure 2A**), with some individual cells missed entirely or only partially captured due to the discontinuous, circular geometry of the Visium spot grid (**Figures 2B, 2B’**). To infer the most prominent cell type per spot, probabilistic cell-type identity was assigned based on the Allen Institute for Brain Science’s scRNAseq reference atlas^20^, yielding a spatial distribution that recapitulated known laminar organization of M1 (**Figure 2C**). CSN-associated markers such as *Fezf2*, *Bcl11b*, *Crym*, and *Rbp4*^10,11^ were broadly expressed across deep-layers (**Figure 2D**). This widespread expression limits their specificity for unambiguous identification of CSNs. To overcome this constraint, we mapped the spatial distribution of tdTomato transcripts (**Figure 2E**) and fluorescent detection of tdTomato protein (**Figure 2F**), revealing a highly specific CSN-enriched signal which we selected for downstream analysis (**Figure 2G**). We confirmed CSN identity using our previously generated scRNAseq data of *bona fide* adult CSNs^9^ (**Figure 2H**). Deconvolution restricted to CSN-enriched spots resolved CSN subtypes, including CSN-IT, -PT, -CT, and -NP populations (**Figures 2I, 2J**) and illustrated the abundance and spatially restricted distribution of CSNs among other cortical cell types (**Figure 2J’**).

**Figure 2.**
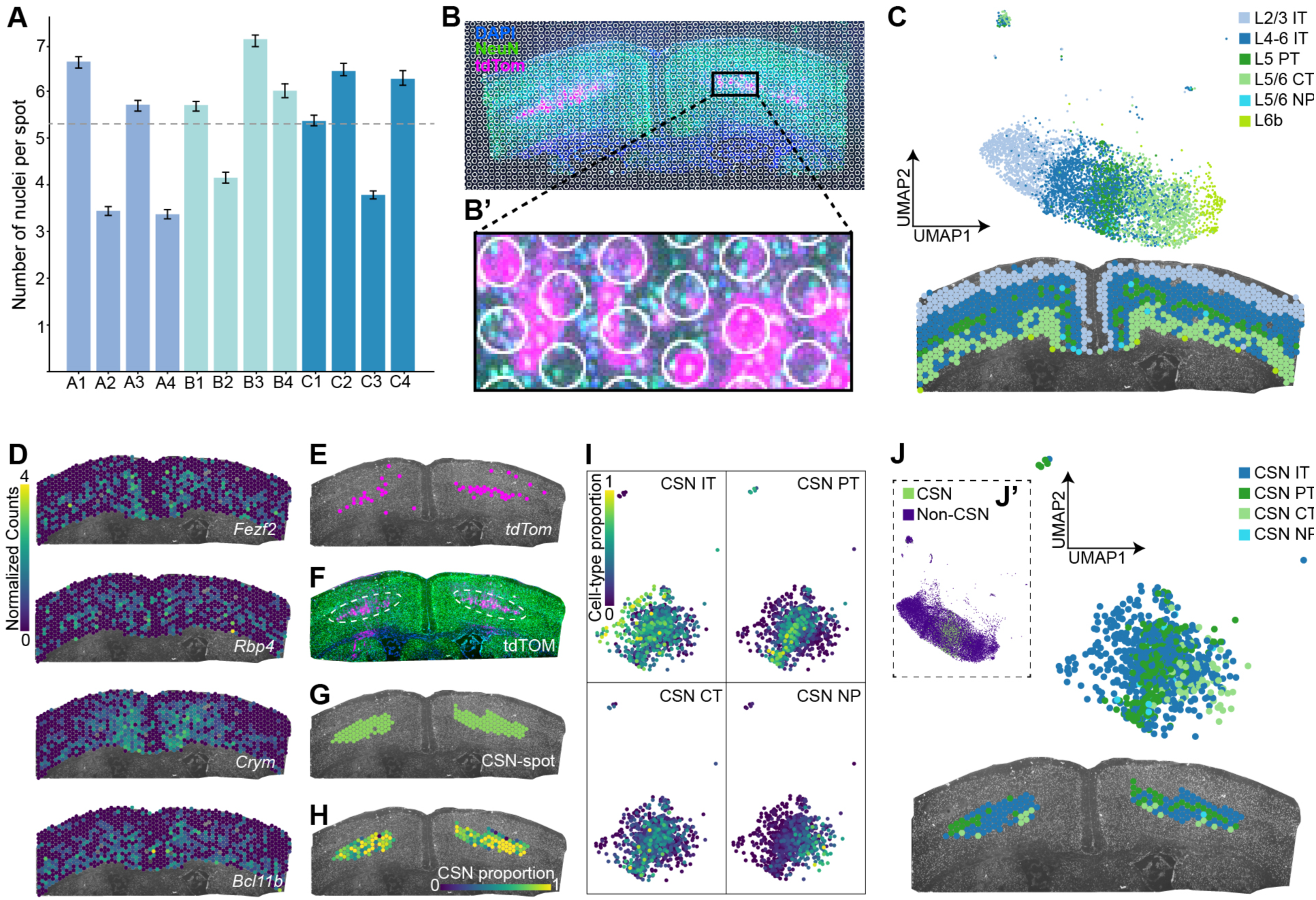
Cell-type deconvolution identified cortical layer and CSN subpopulations. **(A)** Number of nuclei per 55-µm Visium spot. Data represents mean ± SEM; n = 12 mice. Mean across mice = 5.32 (dashed line); SEM = 0.38. **(B)** Visium spot grid overlaid on cortical tissue. **(B′)** Higher-magnification view illustrating spatial resolution. **(C)** Cortical cell-type deconvolution using the Allen Institute M1 reference atlas. Predicted cortical identities are shown in UMAP embedding for all samples space (top) and for a representative tissue section (bottom). IT, intratelencephalic; PT, pyramidal tract; CT, corticothalamic; NP, near-projecting. **(D)** Spatial distribution of canonical layer 5 and CSN marker gene expression, shown as log-normalized counts. **(E)** Spatial localization of tdTomato transcripts. **(F)** Immunofluorescence detection of tdTomato-labeled CSNs. **(G)** Spots of interest selected for downstream analyses. **(H)** Validation of selected CSN spots using a previously generated adult CSN single-cell RNA-seq reference dataset. Estimated CSN enrichment within each spatial spot is shown (0–1). **(I)** Deconvolution of CSN-enriched spots into subpopulations using the Allen Institute M1 reference atlas. Estimated proportions of each CSN subtype per spatial spot are shown in UMAP space (0–1). **(J)** Predicted CSN subtype identities shown in UMAP space (top) and tissue space (bottom). **(J′)** Inset showing CSN-enriched spots relative to all other cortical spots.

### Injured and spared CSNs exhibit distinct transcriptional signatures

Having identified CSN-enriched spots, we examined how injury impacts their transcriptional states by comparing gene expression across intact, injured, and spared conditions (**Figure 3**). Principal component analysis (PCA) revealed separation across conditions, with injured CSNs clearly segregated along PC2, while intact and spared CSNs clustered more closely together (**Figure 3A**). We conducted all biologically meaningful pairwise contrasts between these populations (**Figure 3B**). First, injured vs intact CSNs (purple, X) were compared to isolate the injury-induced transcriptional program. Second, spared vs intact CSNs (blue, S + cX) were compared to capture spared-related changes within the broader injury context (cX) experienced by the animal. Third, to resolve the spared-specific program, we compared spared vs injured CSNs (cyan, S / X) within uPyX animals. Fourth, we performed a control comparison between hemispheres within intact animals (grey, I) to account for background variability. To assess the relationships among these contrasts, we projected each differential expression result into a two-dimensional similarity space defined by an “injury axis” (X) and a “spared axis” (S / X). The spared versus intact contrast (S + cX) occupied a distinct quadrant, suggesting a mixed transcriptional signature (**Figure 3C**).

**Figure 3.**
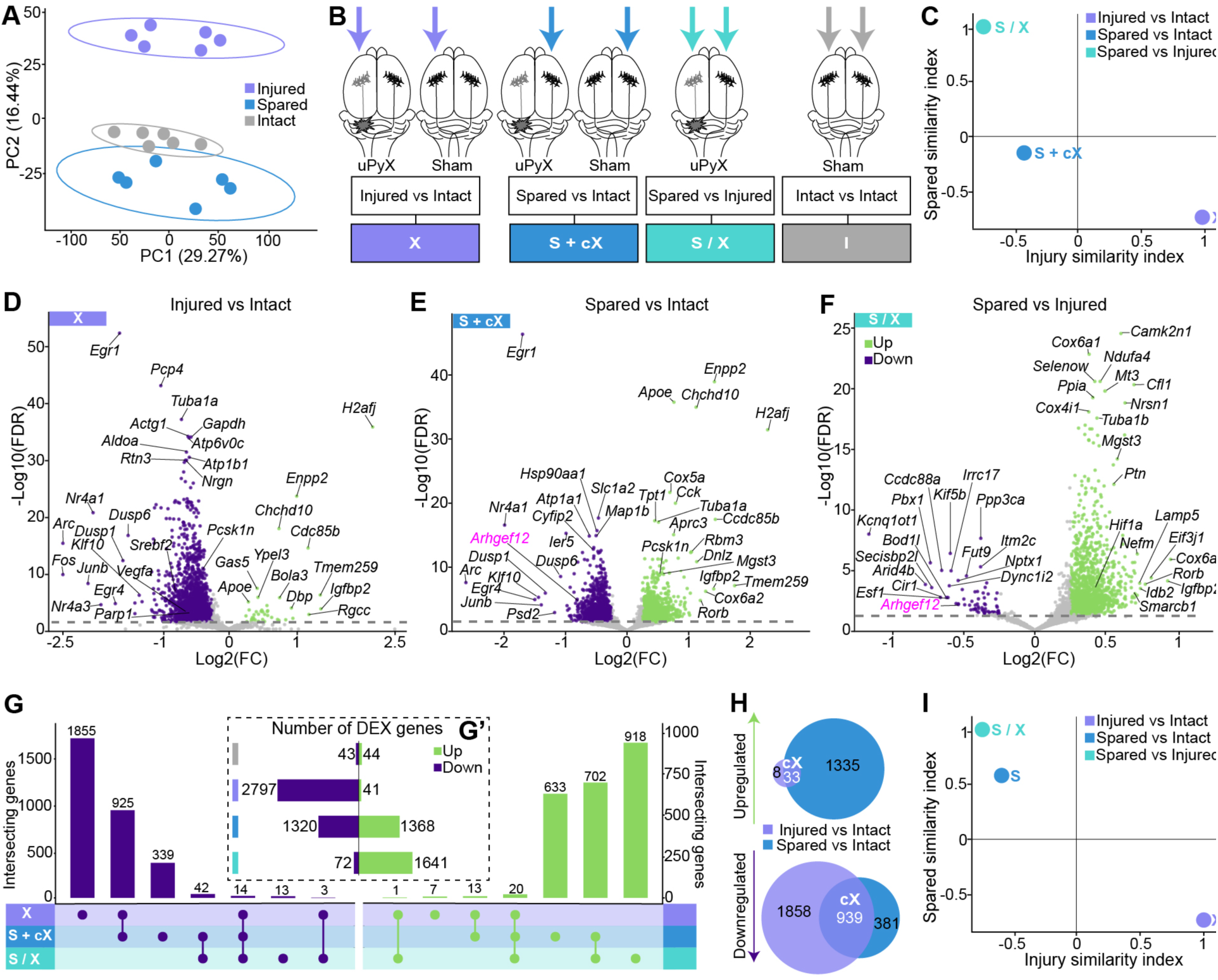
Injured and spared CSNs exhibit distinct transcriptional signatures. **(A)** PCA of CSNs pseudobulk-normalized gene expression from CSN-enriched spots per animal. n = 6 mice per group. **(B)** Schematic of differential expression contrasts performed. **(C)** Two-dimensional similarity-space projection of differential expression contrasts along injury and spared transcriptional axes, using X and S/X as respective anchors. **(D–F)** Volcano plots showing differentially expressed genes for injured vs intact **(D)**, spared vs intact **(E)**, and spared vs injured **(F)** contrasts. **(G)** Intersection of differentially expressed genes among contrasts. **(G′)** Number of significantly upregulated and downregulated genes in each contrast. **(H)** Overlap of differentially expressed genes between injured vs intact (X) and spared vs intact (S + cX) contrasts, highlighting shared injury-context (cX) genes. **(I)** Similarity-space projection of refined spared vs intact (S) signature after computational removal of shared injury-context genes. Differential expression was assessed using the Wald test implemented in PyDESeq2, with Benjamini–Hochberg correction for multiple comparisons. DEGs were defined by padj ≤ 0.05 and |Log2(FC)| ≥ 0.25.

Gene-level differential expression analysis revealed a largely downregulated transcriptome in injured CSNs (**Figure 3D**), while spared CSNs showed both up- and downregulation (**Figure 3E**). There was a substantial overlap of DEGs among the injured vs intact (X) and spared vs intact (S + cX) contrasts (**Figures 3D, 3E**), encompassing transcriptional regulators, activity-responsive genes, and stress-associated phosphatases, including Klf10, Nr4a1, Egr1, Egr4, Arc, Junb, Dusp1, and Dusp6. The spared vs injured (S / X) contrast was more distinct, characterized by a predominantly upregulated transcriptome in spared CSNs and minimal overlap with dysregulated genes from the injured vs intact (X) contrast (**Figure 3F**). We found 2,838 DEGs in the injured vs intact (X, down = 2,797; up = 41), 2,688 DEGs in the spared vs intact (S + cX, down = 1,320; up = 1,368), 1,713 DEGs in spared vs injured (S / X, down = 72; up = 1,641), and only 87 DEGs in the control intact vs intact contrast (I, down = 44; up = 43) (**Figure 3G’**). 972 DEGs were dysregulated in the same direction between the injured vs intact and spared vs intact comparisons (X and S + cX, down = 939; 33 = up) (**Figures 3G, 3H**). These genes were defined as “shared injury context” (cX, **Figure 3H**) and subtracted to isolate a spared-specific signature (S). Projection of the refined signature (S) into the similarity space highlighted its localization within the spared-associated quadrant (**Figure 3I**).

### Functional analysis reveals convergence of spared-specific DEGs on plasticity programs

To characterize the biological processes and regulatory mechanisms associated with each transcriptional signature, we assessed functional pathway and transcription factor enrichment (**Figure 4**). At the pathway level the two spared-specific signatures (S and S / X) were highly correlated to each other (*ρ* = 0.89), and strongly anticorrelated (*ρ* = -0.85, -0.91) with the injury signature (X, **Figure 4A**). The injury profile (X) was characterized by strong pathway inhibition, while both spared-specific profiles (*S* and S */* X) showed widespread activation (**Figure 4B**). Specifically, Ingenuity Pathway Analysis (IPA) ^21^ *Canonical Pathways* related to metabolic support and biosynthesis (Oxidative Phosphorylation, INPP, Cholesterol Biosynthesis), transcriptional and translational machinery (mRNA Polyadenylation, RUNX1/2, Protein Folding), and transport (COPI/II, ABC, Mitochondrial Protein Import), were predicted to be activated in spared CSNs. Pathways linked to structural remodeling and connectivity, such as growth factors signaling (TRKA, IGF1, FGFR2), axon guidance (Ephrin, ROBO, Netrin, L1CAM, Reelin), cell adhesion (CDH1, Cadherin Activation of STAT3, Remodeling of Adherent Junctions), as well as cytoskeletal dynamics (Actin Motility, ILK, ARP-WASP Complex), also had high activation scores (**Figure 4C**). Additionally, Infection of Cells, Macromolecule Metabolism, Cell Survival, Organization of Cytoskeleton, and Microtubule Dynamics were among the most enriched *Diseases and Biological Functions* in spared CSNs, which further supports a growth-permissive, pro-plasticity state. (**Figure 4D**). In contrast, injured CSNs exhibited selective predicted activation of stress and cell death *Canonical Pathways* (Mitochondrial Dysfunction, Apoptosis, Parkinson’s Signaling), coupled with Growth Failure, and Motor Dysfunction as the most enriched *Diseases and Biological Functions* (**Figures 4C, 4D**).

**Figure 4.**
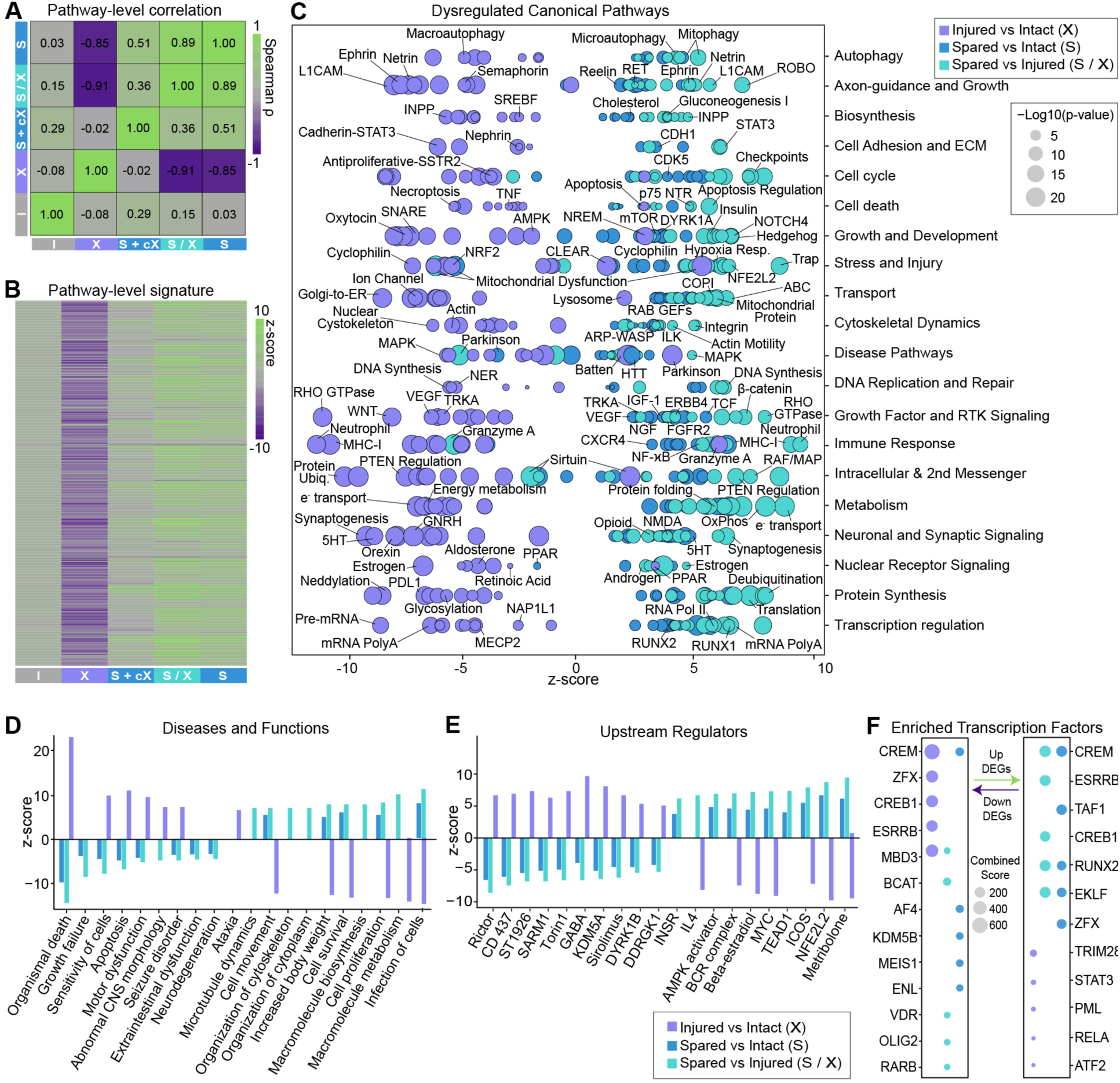
Functional analysis reveals convergence of spared-specific DEGs on plasticity programs. **(A)** Weighted Spearman correlation matrix of *Canonical Pathways* activation z-scores across all biological comparisons. **(B)** Heatmap of the 500 *Canonical Pathways* with the highest absolute activation z-scores across transcriptional signatures. **(C)** Top 10 enriched *Canonical Pathways* per biological category and signature, ranked by enrichment significance. **(D)** Selected 10 predicted activated and 10 predicted inhibited *Diseases and Biological Functions* across signatures, based on activation z-scores. **(E)** Selected of 10 predicted activated and 10 predicted inhibited *Upstream Regulators* across signatures, based on activation z-scores. **(F)** Top 5 enriched transcription factors per signature and direction, ranked by Enrichr Combined Score. (**A-F**) DEGs (padj ≤ 0.05, |Log2(FC)| ≥ 0.25) were used for all functional analyses in IPA and Enrichr. Significance thresholds were *p* ≤ 0.05 for IPA and padj ≤ 0.05 for Enrichr.

IPA *Upstream Regulators* analysis predicted activation of the metabolic and growth regulator MYC^22^, antioxidant response regulator NFE2L2^5^, and anti-inflammatory cytokine IL-4^23^ in spared CSNs. Growth-associated hormonal signaling, including β-estradiol^24^ and androgen receptor signaling^25,26^ (metribolone), and the YAP/TAZ-associated transcription factor TEAD1^27^ were also predicted to be activated. These predictions were accompanied by inhibition of the mTOR pathway^28^ antagonists sirolimus (rapamycin) and Torin1, consistent with enhanced growth-associated signaling (**Figure 4E**). Several other predicted regulatory changes in spared CSNs suggested reduced constraints on neuronal growth and plasticity. These included suppression of the chromatin-associated repressor KDM5A, a chromatin-associated H3K4 demethylase whose degradation promotes neurite outgrowth^29^, the axon degeneration-associated NAD+ hydrolase SARM1^30,31^, and modulation of GABAergic inhibitory neurotransmission, a key regulator of cortical plasticity and circuit reorganization^32^ (**Figure 4E**).

Additionally, transcription factor enrichment analysis (Enrichr^33,34^) predicted activity-dependent and pro-plasticity regulators, CREM^35^, RUNX2^36^, and EKLF^37^ to be enriched in spared CSNs (**Figures 4F**). In contrast, upregulated genes in injured CSNs were enriched for injury- and stress-associated transcriptional regulators, including STAT3^38^, RELA^39^, and ATF2^40^ (**Figure 4F**).

### Network-based prioritization identifies *Arhgef12* as a negative regulator of neurite regeneration

A network-based approach prioritized DEGs common to both spared-specific signatures (S and S / X) and related to axonal growth and repair for functional validation (**Figure 5**). 702 up- and 42 down-regulated DEGs were shared in the spared-specific comparisons (S and S / X) (**Figure 5A**). After filtering for the structural plasticity *Canonical Pathways* significantly enriched across both spared-specific comparisons (e.g., Actin Cytoskeleton Dynamics, Axonal Guidance, Reelin Signaling, and Rho GTPase Signaling, **Figure 5B**), we observed increased expression of genes involved in cytoskeletal remodeling and axonal branching (*Cfl1*, *Tubb4a*, Arp2/3 complex subunits)^41–43^ in spared CSNs. Genes in the RhoA growth-inhibitory signaling cascade (*Rock2*, *Arhgef12*)^44,45^ were consistently among the most downregulated. Since *Arhgef12* was recurrently present in all these pathways, we used IPA Molecule Activity Predictor (MAP) to better understand the functional impact of *Arhgef12* modulation. Downregulation of *Arhgef12* was predicted to increase neuritogenesis and growth of neurites (**Figure 5C**).

**Figure 5.**
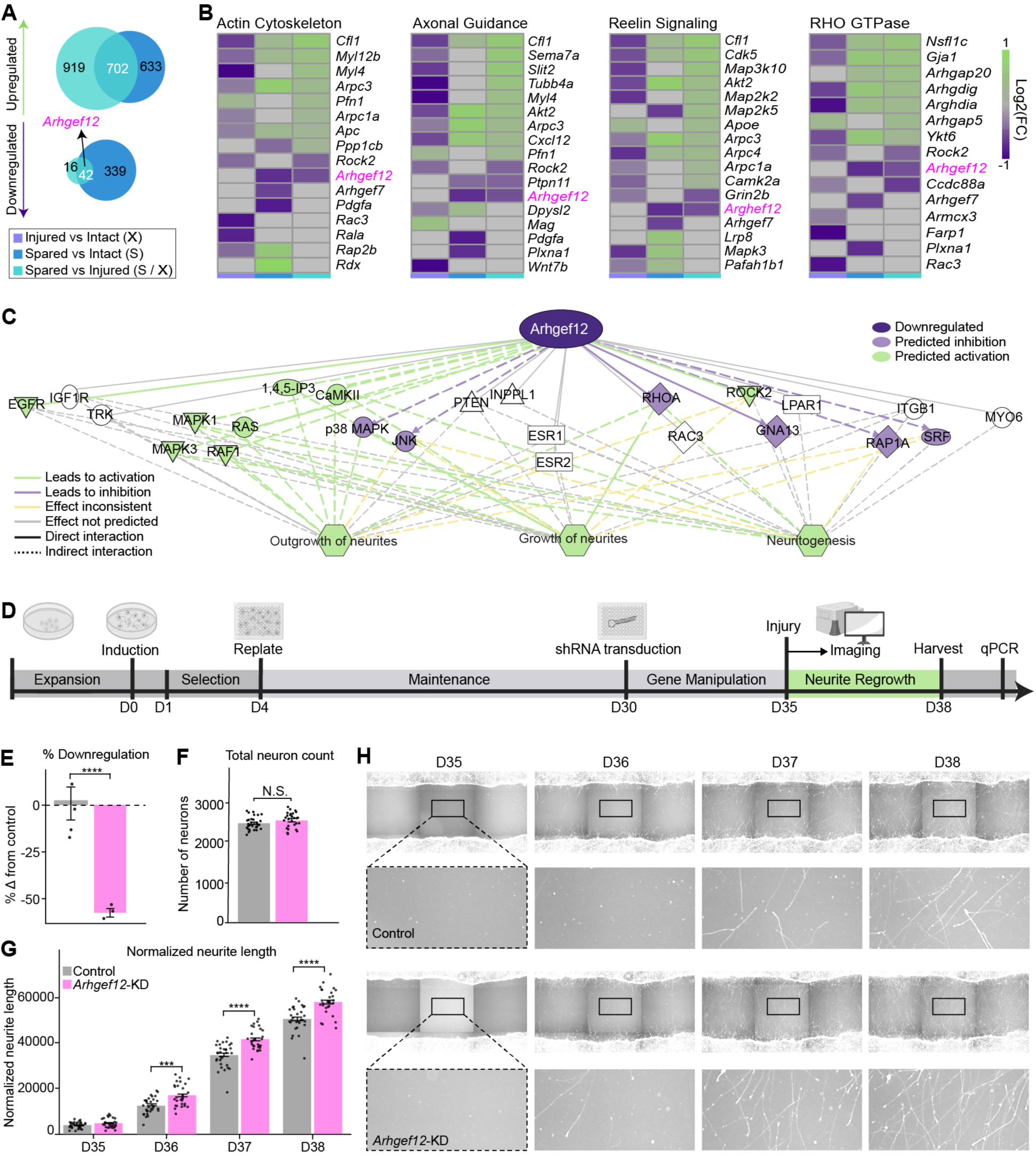
Network-based prioritization identified *Arhgef12* as a negative regulator of neurite regeneration. **(A)** Intersection of differentially expressed genes shared between the two spared-specific comparisons. Significance threshold: padj ≤ 0.05 and Log2(FC) ≥ |0.25| **(B)** Heatmaps showing the top 3 up- and top 3 downregulated genes from each contrast within significantly dysregulated structural plasticity pathways. Actin Cytoskeleton Signaling (*p*: S 2.0 x 10^-8^; S / X 9.6 x 10^-6^; X 3.2 x 10^-20^; X 3.2 x 10^-12^), Axonal Guidance Signaling (*p*: S 2.6 x 10^-10^; S / X 1.1 x 10^-7^; X 3.2 x 10^-20^), Reelin signaling (*p*: S 6.8 x 10^-10^; S / X 1.8 x 10^-7^; X 5.0 x 10^-18^), and Rho GTPase Cycle (*p*: S 1.6 x 10^-11^; S / X 4.6 x 10^-9^; X 1.0 x 10^-19^). **(C)** IPA MAP network illustrating predicted effects of *ARHGEF12* downregulation on neuritogenesis and growth of neurites. **(D)** Experimental workflow and timeline of neurite regeneration assay in hiPSC-derived cortical neurons^93^. **(E)** qPCR quantification of *ARHGEF12* expression relative to the control. **(F)** Total neuron counts at the experimental endpoint. **(G)** Quantification of normalized neurite length with post hoc comparisons between *ARHGEF12*-KD and control at each day following injury. **(H)** Representative images of neurite regeneration following mechanical injury in control (top) and *ARHGEF12*-KD (bottom) NGN2-eGFP-labeled neurons. Insets show higher-magnification views of the regenerating region. Data are presented as mean ± SEM. Neurite regeneration: control (n = 30 wells), *ARHGEF12*-KD (n = 28 wells); qPCR: n = 5 per condition. Statistical significance: ns, not significant; *p* < 0.05 (*), *p* < 0.01 (**), *p* < 0.001 (***), *p* < 0.0001 (****).

To validate whether *ARHGEF12* plays an inhibitory role in axonal growth in a human neuronal model, we conducted a loss-of-function experiment in hiPSC-derived *NGN2*-induced functionally mature glutamatergic neurons^46–48^ (**Figure 5D**). At 30 days *in vitro* (DIV) neurons were transduced with lentiviral shRNA constructs for *ARHGEF12* or a non-targeting control. Validation via qPCR confirmed a 59.6% reduction in *ARHGEF12* expression relative to the control (**Figure 5E**). On DIV 35, neurons underwent a mechanical transection, followed by high-content imaging and automated neurite tracing for 3 days. Critically, total neuron number did not differ between groups (**Figure 5F**), indicating that differences in neurite regeneration were not attributable to neuronal survival or plating density. Quantitative analysis using a linear mixed-effects model revealed a significant increase in normalized neurite length in *ARHGEF12*-KD neurons compared with controls at each day following injury (**Figures 5G, 5H**). Together, these findings provide functional validation of the computational predictions and identify *ARHGEF12* as a negative regulator of neurite regeneration in human neurons.

### Drug perturbation analysis identifies vorinostat as a pharmacological enhancer of neurite regeneration

We next leveraged the spared CSN transcriptional signature to identify candidate pharmacological interventions that could enhance neurite regeneration. A drug perturbation enrichment analysis using the LINCS L1000^49^ database on the 702 commonly upregulated genes across the spared-specific comparisons (S and S / X, **Figure 6A**) identified histone deacetylase (HDAC) inhibitors among the highest-ranking compounds, with vorinostat emerging as the most significant candidate (**Figure 6B**). To evaluate its effects on neurite regeneration, we performed a mechanical transection in hiPSC-derived cortical neurons at DIV35, followed by immediate treatment with 1 μM vorinostat or vehicle (DMSO) (**Figure 6C**). Vorinostat significantly increased normalized neurite length at 2 and 3 days post-injury (**Figure 6D**), without significantly affecting total neuron number (**Figure 6D’**). Representative images confirmed injury consistency and further illustrated increased neurite regrowth following vorinostat treatment (**Figure 6E**). Together, these findings demonstrate that transcriptome-guided drug prioritization can identify pharmacological interventions capable of enhancing neurite regeneration in human neurons.

**Figure 6.**
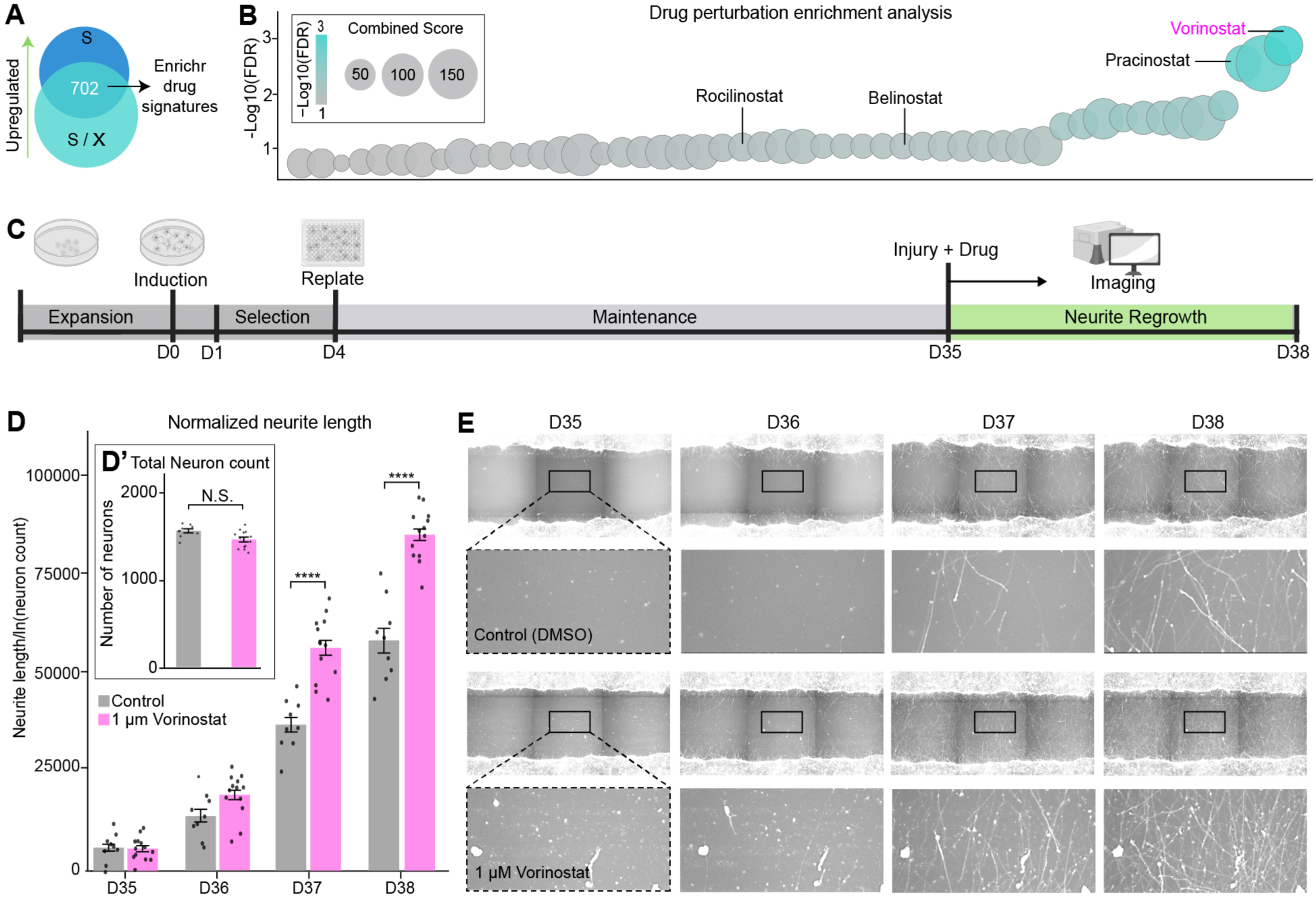
Drug perturbation analysis identifies vorinostat as a pharmacological enhancer of neurite regeneration. **(A)** Drug perturbation enrichment workflow using the 702 upregulated genes in the common spared-specific transcriptional signature as input for the LINCS L1000 Chemical Perturbations up library. **(B)** Top enriched drug perturbation signatures ranked by padj. Bubble size represents the Enrichr combined score. Vorinostat and other HDAC inhibitors are highlighted. **(C)** Experimental workflow and timeline of the pharmacological neurite regeneration assay^93^). **(D)** Pairwise comparisons between vorinostat and vehicle control for normalized neurite length at each day following injury. **(D′)** Total neuron counts at the experimental endpoint. **(E)** Representative images of neurite regeneration following treatment with vehicle (DMSO) or 1 μM vorinostat in NGN2-eGFP-labeled neurons. Insets show higher-magnification views of the regenerating region. Data are presented as mean ± SEM. Neurite regeneration: Control (n = 9 wells), 1 μM vorinostat (n = 13 wells). Statistical significance: ns, not significant; *p* < 0.05 (*), *p* < 0.01 (**), *p* < 0.001 (***), *p* < 0.0001 (****).

## Discussion

In this study we integrated retrograde labeling, spatial transcriptomics, and unilateral injury (uPyX) to profile injured and spared CSNs in primary motor cortex (M1) (**Figure 1**). Robust capture of transcripts, conserved tissue morphology (**Figure S1**), and integration with scRNAseq datasets enabled identification of cortical layers and CSN-enriched spots from known subpopulations^9–11^, including IT, PT, CT, and NP (**Figures 2**). Differential expression and pathway analysis of CSN-enriched spots revealed distinct and opposing transcriptional states in injured and spared populations (**Figures 3, 4**). Injured neurons exhibited a largely downregulated profile, enriched for cell death, motor dysfunction, and metabolic imbalance. In contrast, spared neurons exhibited a predominantly upregulated profile characterized by activation of immune, survival, energy production, translation, protein synthesis, and cytoskeletal remodeling programs. Network-based prioritization of genes dysregulated in spared neurons identified *ARHGEF12* as a candidate negative regulator of neurite regeneration (**Figure 5**), whereas transcriptome-guided drug perturbation analysis identified vorinostat as a compound predicted to engage components of the pro-plasticity signature (**Figure 6**). Consistent with these predictions, *ARHGEF12* suppression and vorinostat treatment independently enhanced neurite regeneration in hiPSC-neurons.

### Distinct intrinsic states shape the transcriptional responses of injured and spared CSNs to common injury cues

The main distinction between injured and spared CSNs is the loss of the primary descending spinal axon in the injured population. Axotomy removes the neuron’s largest cellular compartment, eliminating the extensive corticospinal projection and the continuous biosynthetic and energetic investment required for its maintenance. Consistent with this structural change, injured CSNs exhibited widespread inhibition of ATP production, metabolic regulation, protein synthesis and transport pathways (**Figure 4C**). Additionally, injured CSNs remain embedded within cortical and thalamocortical networks that could convey activity-dependent or trophic signals after distal axotomy^50,51^. However, the loss of their principal motor output creates an imbalance between preserved synaptic input and functional output. Together, these structural, metabolic, and activity-dependent changes may fundamentally reshape the intrinsic state of injured CSNs.

In contrast, spared CSNs retain intact connectivity, preserving both synaptic input and functional output. Despite being anatomically uninjured, they are exposed to altered neuronal activity arising from injury-induced network reorganization, including interhemispheric signaling^52,53^, as well as extrinsic cues associated with the injury response^54^. Following injury, activation of resident glia, infiltration of peripheral immune cells, and the release of cytokines and chemokines initiate a signaling cascade that extends throughout the CNS^55^. In addition to promoting debris clearance and wound healing, this immune response can induce transcriptional reprogramming in both local and distal cells^56,57^. Consistent with this concept, immune pathways were among the most enriched biological programs in spared CSNs (**Figures 4C, 4D**). Yet, unlike injured CSNs, the preserved structural integrity and potentially increased network activity in spared CSNs likely underlies their coordinated upregulation of metabolic, survival, translational, and cytoskeletal remodeling pathways in response to injury-associated signals (**Figures 4C-4F**). Together, these findings suggest that depending on their intrinsic states, common extrinsic cues may contribute to degeneration in injured neurons while simultaneously priming spared CSNs to activate adaptive plasticity programs.

### Spared CSNs integrate metabolic, structural, and survival programs to support plasticity

Spared CSNs exhibited a highly permissive transcriptional state characterized by coordinated activation of biological programs that support not only neuronal survival but also structural and functional reorganization after injury. If spared CSNs receive increased excitatory drive from contralateral cortical circuits following unilateral injury, the initial increase in synaptic output would likely rely on pre-existing cellular machinery. Consistent with this concept, GABA Signaling was among the most de-enriched *Upstream Regulators* in spared CSNs, whereas the activity-dependent transcription factor CREM^35^ and the Synaptogenesis pathway were among the most enriched (**Figures 4C-4F**). Sustained structural remodeling, however, requires substantial energy production and biosynthetic capacity, reflected in the predicted activation of the metabolic regulator MYC^22^ and multiple pathways involved in metabolism, transcription, protein synthesis, and intracellular trafficking (**Figures 4C, 4E**). Importantly, these changes were accompanied by enrichment of pathways directly involved in axon growth and structural remodeling, including Netrin, Ephrin, Semaphorin, SLIT–ROBO, and L1CAM signaling, together with pathways regulating actin cytoskeleton dynamics, RHO-mediated Motility, ARP–WASP-dependent Actin Nucleation, and Integrin–Cytoskeleton Interactions (**Figure 4C**). In parallel, activation of pathways and regulators associated with immune modulation, antioxidant defense, and neuronal survival, including NFE2L2^5^, IL-4^23,58^, STAT3^38^, and mTOR^28^ (**Figures 4C, 4E**), may represent complementary mechanisms to preserve cellular homeostasis and neuronal integrity. Similar activation of metabolic and antioxidant programs has been previously identified in regenerating CSNs following *Pten/Socs3* deletion^5^, supporting the possibility that the spared CSN transcriptional signature identified here reflects molecular mechanisms associated with axonal repair. Together, these transcriptional programs appear to function as components of a unified adaptive response, aimed at protecting and strengthening existing circuits, while building new connections after injury.

### Endogenous plasticity programs are detectable even if insufficient to drive functional recovery

Despite the robust pathway-level pro-plasticity signature observed in spared CSNs, spontaneous anatomical and functional recovery after injury is limited^59,60^. Several factors may contribute to the disconnect between the observed transcriptional response and the biological outcome. First, the magnitude of individual gene expression changes remained relatively modest, with most differentially expressed genes in both spared and injured CSNs exhibiting Log2(FC) < 1 (**Figures 3D-3F**). Although our previous study primarily investigated sprouting CSNs in *Ngr1*-null mice, the accompanying wild-type comparison between spared and injured CSNs after uPyX also identified minimal dysregulation^4^. Similarly, a separate study of injured CSNs reported limited transcriptional changes one week after cervical and thoracic spinal cord injury^7^. Our injured CSNs recapitulated the downregulation of activity-dependent genes including Egr1, Egr4, Arc, Nr4a1, and Junb (**Figure 3D**), together with enrichment of injury-responsive transcription factors such as ATF2 (**Figure 4F**). These concordant findings support the biological identity and relevance of the molecular signatures of injured and spared CSNs, while reinforcing the concept that the magnitude of these endogenous responses to distal axotomy may be insufficient to drive substantial functional recovery.

Second, the endogenous pro-plasticity state may be inherently transient. Consistent with this idea, previous characterization of layer V neurons showed that injury alone can induce an early regenerative transcriptome characterized by reversion toward an embryonic-like state^6^. Our spared CSNs exhibited transcriptional features consistent with an immature, growth-permissive state, including increased HTT upregulation (**Figures 4C, 4D**). However, in that study in the absence of stem cell grafts, this transcriptional program progressively declined 10 days post lesion. Likewise, we previously demonstrated that untreated mice undergo minimal anatomical reorganization and spontaneous recovery beyond modest initial improvements during the first week post-injury, while motor rehabilitation significantly enhanced corticospinal remodeling and continued functional recovery^61^. Collectively, these findings suggest that intrinsic plasticity programs may require external reinforcement to be sustained.

Third, our analysis suggests that the endogenous pro-plasticity response of spared CSNs is constrained by concurrent injury-associated signaling (**Figures 3E, 3H**). Although computational removal of shared injury-context genes revealed a robust pro-plasticity transcriptional signature, both programs coexist within spared neurons *in vivo*. Consequently, injury-associated transcriptional responses can counterbalance activation of plasticity-related pathways, producing a more muted intrinsic growth state (**Figure 4B**). Importantly, these intrinsic constraints are compounded by the inhibitory extracellular environment that develops after CNS injury. Reactive gliosis and scar formation generate a complex lesion environment in which chondroitin sulfate proteoglycans, myelin-associated inhibitors, and persistent neuroinflammation can restrict axonal growth and structural remodeling^62–64^. These observations support a model in which spontaneous corticospinal plasticity is constrained at multiple levels: magnitude, duration, and concurrent intrinsic and extrinsic injury-inhibitory signals.

### An integrative framework for target- and state-based therapeutic discovery

Our integrative discovery pipeline demonstrates how transcriptional profiling of spared CSNs can be leveraged to identify both intrinsic genetic regulators and pharmacological interventions that promote neurite regeneration. Our gene prioritization strategy included refinement at multiple levels: isolation of the spared-specific signature, overlapping gene lists from between and within animal comparisons, recurrence of candidates at the pathway-level, and *in silico* predictions of their involvement in neurite growth. Through this approach, *Arhgef12* emerged as a central negative regulator of neurite regeneration. Although *Arhgef12* has been primarily characterized as a Rho guanine nucleotide exchange factor that activates RHOA^65–67^, our analyses predicted broader interactions with multiple cytoskeletal remodeling pathways. Consistent with these predictions, *ARHGEF12* suppression significantly enhanced neurite regeneration in hiPSC-neurons, establishing *ARHGEF12* as a previously unrecognized regulator of regenerative growth.

In parallel, unbiased drug perturbation analysis^33,34,49^ identified histone deacetylase inhibitors as a convergent pharmacological class associated with the spared-CSN transcriptional program. Vorinostat was the highest-ranked compound, while the independent enrichment of pracinostat, belinostat and rocilinostat demonstrated convergence on HDAC inhibition as a pharmacological class rather than an isolated compound-signature match. HDAC inhibitors have shown beneficial effects in experimental stroke and spinal cord injury, where their actions were linked not only to neuroprotection and modulation of inflammatory and oxidative signaling^68–70^, but also to enhanced neurite outgrowth and axon regeneration^71,72^. Vorinostat itself has demonstrated neuroprotective effects following cerebral ischemia and hemorrhage^68,73^, promotes neurite outgrowth in PC12-derived neuronal-like cells^74^, and has been evaluated following traumatic spinal cord injury^75^. However, its potential to directly promote regenerative neurite growth following injury, particularly in human neurons, had not been established. Here, rather than selecting HDAC inhibition *a priori*, our unbiased analysis of spared CSNs independently nominated vorinostat as a compound predicted to engage components of the endogenous pro-plasticity program, while subsequent validation in human neurons demonstrated a direct effect on regenerative growth. These results illustrate how complementary target- and state-based strategies can be identified and validated in a neuronal model that provides initial evidence of their efficacy in a human genetic background.

### Limitations of the study

#### Injury Paradigm

The uPyX model permits selective unilateral targeting of the corticospinal tract at a distal location while preserving the contralateral side and adjacent cortical circuitry^4,76^. Although a uPyX does not fully recapitulate clinically relevant spinal cord injury, it was well suited for transcriptional profiling of a specific neuronal population within a single animal across two distinct states: injured and spared.

#### Spatial Resolution

Spatial transcriptomics enables capture of nuclear and cytoplasmic transcripts directly from the tissue circumventing the need for dissociation and avoiding processing-related gene expression artifacts. This approach combined with retrograde labeling from the spinal cord and reference scRNAseq atlases for deconvolution affords cell type specificity mitigating the limitation of the non-single cell resolution. Moreover, preservation of anatomical architecture can turn resolution constraints into an advantage by including information not only from the target CSNs but also from their adjacent native microenvironment, which can better reflect complex biological states, such as intrinsic gene expression changes of spared neurons coexisting with injury-induced contextual cues. Future studies, at single-cell spatial resolution, thanks to newly released technologies (Visium HD), will allow explicit interrogation of gene expression changes driven by each cell type, including astrocytes and microglia.

#### Temporal Resolution

This study focuses on a single time point (D10) after injury. We selected D10 because endogenous gene expression changes after spinal cord injury have been reported to persist through this stage^6,7^. In addition, we previously observed the onset of spontaneous functional recovery between D7 and D14 after injury^61^, suggesting that transcriptional programs driving circuit reorganization are likely initiated before or during this period. Consistent with this timeline, we also found that motor rehabilitation promotes anatomical reorganization and significantly enhances functional recovery^61^. Future studies will extend these findings by profiling rehabilitation-induced gene expression across multiple post-injury time points to better define the temporal sequence of molecular events underlying spontaneous and rehabilitation-induced plasticity.

#### Human model

To focus on neuron cell-autonomous response to injury, this study tested functionally mature human glutamatergic neurons^46,48^ cultured in the absence of astrocytes, oligodendrocytes, and microglia, support cells that might otherwise impact axon regeneration^62^. Likewise, our assay was conducted in a growth factor depleted state to maximize signal and resolve effects of genetic and pharmacological perturbations, which can alter baseline cell signaling contexts.

In summary, we found that spared CSNs mount an endogenous pro-plasticity response after unilateral injury. This transcriptional signature can be leveraged to identify target and state-based repair strategies such as *ARHGEF12* and vorinostat. Whereas ARHEGF12 suppression engages a defined target linked to RHOA and cytoskeletal remodeling, vorinostat acts at the state level across transcriptional programs associated with the pro-plasticity response. Their independent effects in human cortical neurons support the biological relevance of the spared-CSN transcriptional signature. More broadly, these findings establish a scalable framework integrating spatial transcriptomic discovery with functional screening in human neurons to translate endogenous responses to CNS injury into candidate strategies for neural repair.

## Resource Availability

### Lead contact

Requests for further information and resources should be directed to and will be fulfilled by the lead contact William B J Cafferty.

### Materials availability

This study did not generate new unique reagents.

### Data and code availability

Spatial transcriptomics data have been deposited in the Gene Expression Omnibus (GEO) under accession number (pending). All original code and supporting analysis files have been deposited in Dryad (DOI: pending). These resources will be made publicly available upon publication. Any additional information required to reanalyze the data reported in this paper is available from the lead contact upon request.

## Acknowledgments

This work was supported by grants from the NIH (R01NS121026 and R21NS139481 to W.B.J.C, and R01ES033630, RM1MH132648 to K.J.B), Wings for Life (2023-066 to W.B.J.C), NINDS T32 (NS007224 to M.M.), F31NS143390 (to M.M.), Gruber Science Fellowship (to M.M).

## Author contributions

M.M., W.B.J.C., and K.J.B., conceptualization and experimental design. N.G. and W.B.J.C., surgical procedures. M.M., spatial transcriptomics tissue processing. C.O. and M.M., bioinformatic analysis. M.M. and E.Y., stem cell experiments. M.P.J.D., high-content imaging training. J.M., N.B., and M.P.J.D., stem cell experimental guidance. M.M., data analysis, visualization, and writing. M.M., K.J.B., and W.B.J.C., manuscript editing. K.J.B. and W.B.J.C., supervision and funding acquisition. All authors read and approved the manuscript.

## Declaration of generative ai and ai-assisted technologies in the writing process

During the preparation of this manuscript, the authors used ChatGPT based on GPT-5.5 to improve clarity and readability. After using this tool, the authors reviewed and edited the suggestions and take full responsibility for the content of the publication.

## Supplemental information

**Figure S1.**
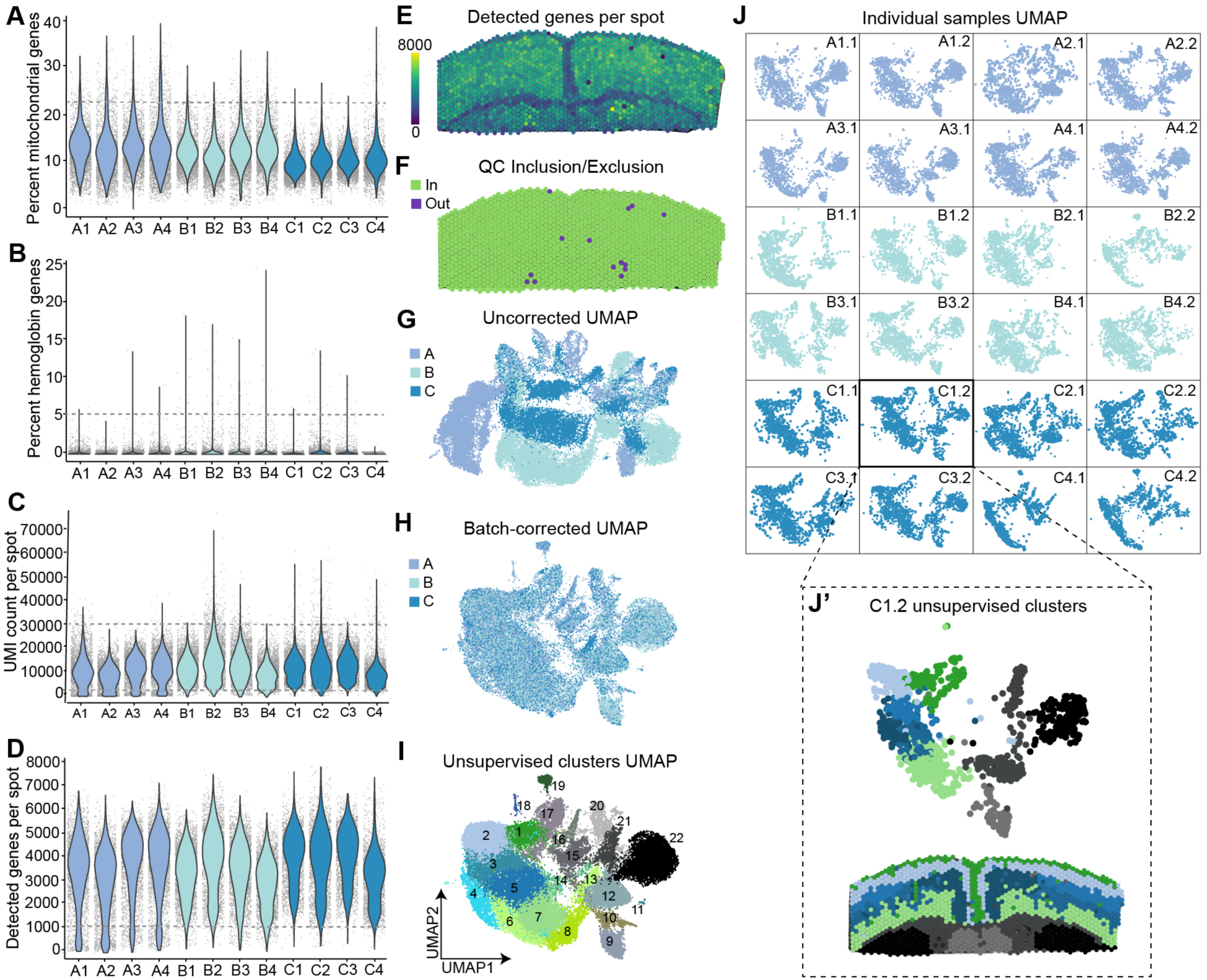
Anatomically resolved gene expression is conserved across biological replicates. (A–D) Quality-control metrics across all Visium capture areas. **(A)** Percent mitochondrial reads, mean = 12.32%. **(B)** Percent hemoglobin reads, mean = 0.10%. **(C)** UMI counts per spot, mean = 11,230.24. **(D)** Detected genes per spot, mean = 3,583.04. Dashed lines represent cutoff thresholds. **(E)** Spatial distribution of detected genes across a representative tissue section. **(F)** Spots included (green) or excluded (purple) following quality-control filtering. Inclusion thresholds: < 22.5% mitochondrial counts, < 2.5% hemoglobin counts, > 1,000 genes, and between 1,000 and 30,000 UMIs. **(G)** UMAP before batch correction. **(H)** UMAP following Harmony batch correction. **(I)** Unsupervised Leiden clustering for all samples combined. **(J)** UMAP distribution across individual tissue sections. **(J′)** UMAP (top) and spatial localization (bottom) of unsupervised clusters for representative tissue section. Data are shown for n = 12 mice, represented by 24 tissue sections. Individual Visium spots are shown as individual data points.

## Methods

### Experimental model details

#### Animals

Adult C57BL/6J mice (The Jackson Laboratory) (n = 12; 10 females and 2 males) were randomly assigned to receive unilateral pyramidotomy (uPyX; n = 6) or sham surgery (n = 6). Animals were housed under standard laboratory conditions with *ad libitum* access to food and water and maintained on a 12 h light/dark cycle. All procedures were approved by the Yale University Institutional Animal Care and Use Committee (IACUC) and performed in accordance with NIH guidelines.

#### Human Induced Pluripotent Stem Cells (hiPSCs)

Human induced pluripotent stem cell (hiPSC) line NSB2607 clone 2 (2607-2; karyotype XY) was previously established and characterized^77^ and is available from the NIMH Repository (study 160). Stem cell-derived NGN2-accelerated neural progenitor cells (iNPCs^87,88^) were previously derived and validated. De-identified repository-available hiPSCs were deemed not human subjects research by the Yale School of Medicine Institutional Review Board. Cell cultures were routinely tested and confirmed negative for mycoplasma contamination.

### Method details

#### Retrograde labeling

Adult C57BL/6J mice underwent bilateral retrograde labeling of corticospinal neurons (CSNs) by intraspinal injection of AAVrg-CAG-tdTomato. Mice were anesthetized by intraperitoneal injection of ketamine (100 mg/kg) and xylazine (10 mg/kg). The adequacy of anesthesia was confirmed by the absence of corneal and toe-pinch reflexes. Ophthalmic lubricant was applied to both eyes to prevent corneal drying during surgery. Mice were secured in a stereotaxic frame using atraumatic ear bars, and the surgical site was shaved, scrubbed with betadine, and swabbed with 70% ethanol. A bilateral laminectomy was performed at either the cervical (C6/C7) or lumbar (L4/L5) spinal cord. A pulled glass capillary attached to a 5 μL Hamilton syringe mounted on a Micro4 infusion pump (World Precision Instruments) was positioned using stereotaxic guidance and inserted to a depth of 500 μm and 600 μm lateral to the midline. 30s after insertion, 100 nL of AAVrg-CAG-tdTomato was infused over 2 min at each injection site. The capillary was left in place for an additional 30s before being slowly withdrawn to minimize reflux. This procedure was repeated 7x at evenly spaced injection sites. The surgical site was closed with Vicryl absorbable sutures, and animals received postoperative analgesia (buprenorphine, 0.05 mg/kg, subcutaneous) and antibiotics (ampicillin, 100 mg/kg, subcutaneous) for 2 days after surgery. Mice recovered for 14 days to allow retrograde transport to the cortex and robust tdTomato expression prior to uPyX or sham surgery.

#### Unilateral pyramidotomy (uPyX)

Mice were anesthetized as above. Mice were positioned in the supine position, ophthalmic lubricant applied to both eyes and the neck area shaved and scrubbed with betadine and alcohol. A ventral midline incision was made to the left of the trachea. Blunt dissection was performed to expose the occipital bone at the base of the skull. The occipital bone was carefully removed on the left side of the basilar artery using blunt Dumont #2 forceps to expose the medullary pyramids. The dura mater was pierced with a 30-gauge needle and resected. The left medullary pyramid was transected unilaterally to a depth of 0.25 mm using fine Dumont #5 forceps, whereas sham-operated animals underwent identical surgical exposure without transection of the pyramid. No internal sutures were placed, and the skin incision was closed using monofilament sutures. Animals received postoperative analgesia (Buprenorphine, 0.05mg/kg subcutaneously) and antibiotics (Ampicillin, 100mg/kg subcutaneously) for 2 days post-surgery and were monitored daily throughout recovery.

#### Tissue preparation for Visium spatial transcriptomics

Ten days following unilateral pyramidotomy (uPyX) or sham surgery, mice were deeply anesthetized with isoflurane until loss of the toe-pinch reflex and transcardially perfused with ice-cold 0.9% saline supplemented with heparin. Brains were rapidly removed and immediately transferred to ice-cold saline for dissection. Using sagittal and coronal brain matrices, the forebrain was trimmed by making a coronal cut 3 mm caudal to the olfactory bulb followed by sagittal cuts 3 mm lateral to the midline on each hemisphere, thereby isolating a tissue block containing the primary motor cortex (M1). Tissue blocks were embedded in Tissue-Plus O.C.T. Compound, snap frozen on powdered dry ice, and stored at −80°C until cryosectioning. Prior to sectioning, tissue blocks were equilibrated in a cryostat maintained at −12°C for 30 min. Blocks were trimmed to the level of M1 using tdTomato fluorescence as a guide and scored with a razor blade below the corpus callosum to enable placement of two sections within a single 6.5 × 6.5 mm Visium capture area. The two M1 coronal sections (10 μm) from each animal were collected onto a room temperature Visium Spatial Gene Expression v1 Slide ^12,89^. Sections from uPyX and sham mice were distributed equally on 3 Visium slides (Slides A-C) to control technical variability, with two sections included per animal within each capture area. Immediately after collection, sections were fixed in pre-chilled 100% methanol for 30 min at −20°C.

#### Immunofluorescence

Immediately following fixation, tissue sections were processed for immunofluorescence according to the ‘Methanol Fixation, Immunofluorescence Staining & Imaging for Visium Spatial Protocols’ (CG000312). All solutions, including blocking buffer, primary and secondary antibody solutions, wash buffer, SSC, and mounting medium, were prepared according to the manufacturer’s instructions. To preserve RNA integrity, the recommended blocking and antibody incubation times were reduced. The primary antibody solution contained rabbit anti-mCherry antibody (1:1000), and the secondary antibody solution contained mouse anti-NeuN monoclonal antibody (3A4C1), CoraLite Plus 488 (1:100), goat anti-rabbit IgG Alexa Fluor 546 (1:1500), and DAPI (1:3000). All incubations were performed at room temperature. Briefly, sections were incubated in blocking buffer for 5 min, followed by incubation in primary antibody solution for 10 min. After 5x washes, sections were incubated in secondary antibody solution for 10 min, washed an additional 5x, immersed 20X in SSC solution, and cover slipped.

#### Microscopy

Whole-slide fluorescence images were acquired immediately following immunostaining using a Leica upright epifluorescence microscope DM5500 equipped with a 10X objective, following the Visium imaging workflow (CG000241). Identical acquisition settings were used for all slides to ensure consistent fluorescence intensity across samples. Following imaging, slides were stored overnight at 4°C prior to tissue permeabilization and library preparation.

#### RNA quality assessment

Adjacent M1 sections were collected during cryosectioning into 1.5-mL microcentrifuge tubes for RNA quality assessment. Total RNA was isolated using the RNeasy Mini Kit (Qiagen) according to the manufacturer’s instructions. RNA concentration and purity were initially assessed using a NanoDrop spectrophotometer, and RNA integrity was subsequently determined by the Yale Center for Genome Analysis (YCGA) RNA Quality Control Service using an Agilent 2200 TapeStation to obtain RNA Integrity Number (RIN) values.

#### Permeabilization and barcoded cDNA generation

All tissue permeabilization and barcoded cDNA generation were performed according to the Visium Spatial Gene Expression User Guide (CG000239). The optimal tissue permeabilization time for mouse M1 was determined prior to processing experimental samples using the Visium Spatial Tissue Optimization Kit (CG000238). Based on these results, experimental tissue sections were permeabilized for 3 min. Total polyadenylated mRNA was captured by the oligonucleotides within each Visium capture area containing a Poly(Dt) region, spatial barcodes, unique molecular identifiers (UMIs), and TruSeq Read 1 sequencing adapters. Reverse transcription was performed in situ to generate spatially barcoded first-strand cDNA, followed by second-strand cDNA synthesis, denaturation, retrieval of the barcoded free cDNA strand and amplification prior to submission to the YCGA.

#### Library preparation and sequencing

Sequencing libraries were prepared by the YCGA according to the Visium Spatial Gene Expression User Guide (CG000239). Library construction included cDNA cleanup, quality control, fragmentation, end repair, A-tailing, double-sided SPRIselect size selection, adaptor ligation, post-ligation cleanup, sample index PCR, post-PCR double-sided size selection, and final library quality control prior to sequencing. Libraries were sequenced on an Illumina NovaSeq 6000 platform using paired-end 150-bp sequencing, targeting 177.5 million reads per Visium capture area.

#### Spatial sequencing data preprocessing

Raw sequencing data were processed using Space Ranger v1.3.1 (10x Genomics), which aligned fluorescence tissue images and fiducial marker frames to the Visium capture areas and sequencing reads to a custom mm10_tdTomato reference transcriptome to generate spot-by-gene count matrices for each capture area. Count matrices were imported into the scverse Python environment (Python v3.12.9, Scanpy v1.11.5, custom fork of Squidpy based on v1.6.2) for downstream preprocessing and quality control. For each spot, the total UMI counts, number of detected genes, and percentages of mitochondrial and hemoglobin transcripts were calculated. Spots were retained for downstream analyses if they contained <22.5% mitochondrial transcripts, <2.5% hemoglobin transcripts, >1,000 detected genes, and between 1,000 and 30,000 UMIs. Principal component analysis (PCA) was performed on the filtered dataset. To minimize technical variation between Visium slides, batch correction was performed using Harmonypy (v0.0.10), using slide identity as the batch variable. Batch-corrected principal components were used to construct a nearest-neighbor graph, which was subsequently used for UMAP visualization and Leiden clustering. To isolate cortical tissue for downstream analyses, Leiden clusters from each tissue section were projected onto their corresponding spatial coordinates. Based on their spatial localization, clusters corresponding to the cerebral cortex overlying the corpus callosum were identified and recorded for each tissue section. Only spots within the cortical region of interest were retained for subsequent analyses.

#### Cell type deconvolution

Nuclei abundance per Visium spot was estimated from DAPI fluorescence using a custom ImageJ plugin that quantified DAPI-positive pixel area and normalized it to a predefined nuclear area. Cell-type composition of cortical Visium spots was estimated using Stereoscope (scvi-tools) (v1.4.1) with a previously published Allen Institute primary motor cortex (MOp) single-cell RNA sequencing reference dataset ^20^. Prior to model training, reference cell populations were balanced by randomly subsampling a maximum of 500 cells per subclass, and subclasses represented by a single cell were excluded. Marker genes were identified by between-groups differential expression analysis within the reference dataset and intersected with a list of genes identified as highly variable in the Visium dataset. The RNA model was trained for 300 epochs, followed by training of the spatial model for 3,000 epochs. Estimated cell-type proportions were calculated for every cortical Visium spot and used to annotate the major cellular composition of the cortical region. CSN-enriched spots were identified in Loupe Browser (v6.3.0; 10x Genomics) by integrating anatomical location and tdTomato immunofluorescence. Selected spots were subsequently evaluated using a previously generated adult CSN single-cell RNA sequencing reference dataset ^9^ to confirm enrichment for the CSN transcriptional signature. Following validation, the selected spots were subjected to a second round of Stereoscope deconvolution using the Allen Institute primary motor cortex reference dataset ^20^ to classify CSN subtypes.

#### Assessment of lesion completeness

To assess lesion completeness, PyDESeq2 (v0.5.3)-normalized gene expression matrices from CSN-enriched spots were exported and aggregated into animal-level pseudobulk profiles using mean counts across spots. Prior to principal component analysis (PCA), a linear model was used to regress out slide-specific technical effects, including slide x condition interactions, while preserving the biological condition effect. PCA was performed on the 5,000 most variable genes after centering and scaling the corrected expression matrix. The PCA plot was examined to determine whether samples clustered according to experimental condition and to assess inter-animal variability.

#### Differential expression analysis

Differential expression analysis was performed using PyDESeq2 (v0.5.3) on raw gene count matrices extracted from CSN-enriched spots. Pairwise comparisons between intact left, intact right, spared, and injured CSN-enriched spots were performed for each Visium slide (**Figure 3B**). Raw counts were normalized by estimating sample-specific size factors using the PyDESeq2 median-of-ratios method. Gene-wise dispersions were estimated and negative binomial generalized linear models were fitted for each gene. Differential expression was evaluated using Wald tests. Cook’s distance was calculated to identify influential observations, and count outliers were refitted when appropriate. Independent filtering and Benjamini-Hochberg correction were applied to adjust for multiple testing. Differential expression results were subsequently post-processed in R (v4.3.3) prior to downstream analyses. Genes associated with technical artifacts, including mitochondrial genes, major ribosomal protein gene families, hemoglobin genes, sex chromosome-associated genes, and predicted loci, were removed from each dataset. Results from biological replicates were consolidated into a consensus differential expression table by resolving duplicate gene entries according to the largest |Log2(FC)| in concordant calls and highest significance (lowest padj) in discordant calls. Genes with padj ≤ 0.05 and |Log2(FC)| ≥ 0.25 were considered significantly differentially expressed for downstream analyses and visualization.

#### Computational isolation of a spared-specific transcriptional signature

Consensus differential expression signatures were analyzed in R (v4.3.3). Prior to downstream analyses, genes with a baseMean <2 or not detected across all biological comparisons were excluded, and the resulting data were standardized using gene-wise Z-score normalization. The injured vs intact (X) and spared vs injured (S / X) differential expression contrasts were used as anchors to define injury and spared similarity indices. Cosine similarity scores between each differential expression signature and the anchor signatures were calculated and used to project all biological comparisons onto a two-dimensional injury-spared similarity map. Genes significantly dysregulated in the same direction in both the injured versus intact (X) and spared versus intact (S + cX) comparisons were identified and removed from the spared versus intact comparison to generate a filtered spared-specific signature (S). The refined signature was subsequently reprojected onto the injury-spared similarity map.

#### Functional enrichment analysis

Differentially expressed genes from each comparison were analyzed using Ingenuity Pathway Analysis^21^ (IPA, Qiagen; Content Version 165342674) to identify enriched canonical pathways, diseases and biological functions, and upstream regulators. Enrichment significance and predicted activation state were determined using IPA enrichment statistics and activation z-scores. Canonical pathways were grouped into major biological categories for visualization. To compare functional signatures across biological comparisons, weighted Spearman correlations were calculated from IPA pathway activation z-scores, with pathway weights proportional to enrichment significance (−Log10(p-value), maximum weight = 50). The resulting correlation coefficients were visualized as similarity matrices. Canonical pathways were subsequently visualized as heatmaps and bubble plots, whereas diseases and biological functions and upstream regulators were ranked according to IPA activation z-scores and displayed as bar plots.

#### Transcription factor enrichment analysis

Transcription factor enrichment analysis was performed using Enrichr^33,34^ with the *ChEA 2022* and *ENCODE and ChEA Consensus TFs from ChIP-X* libraries. These libraries identify transcription factors whose experimentally defined target genes are significantly overrepresented among the differentially expressed genes. Upregulated and downregulated genes were analyzed independently for each differential expression signature. Significantly enriched transcription factor terms (padj ≤ 0.05) from both libraries were consolidated by transcription factor, and the highest-ranking transcription factors were used for downstream visualization and comparison across differential expression signatures.

#### Candidate gene prioritization

Candidate genes were prioritized by identifying genes significantly dysregulated in the same direction in both spared-specific signatures (S and S / X). Structural plasticity pathways that were significantly enriched in both spared-specific comparisons were then examined, and the top 3 upregulated and top 3 downregulated genes for each contrast ranked by |Log2(FC)| were selected from each pathway. Genes recurrently represented across multiple pathways were further evaluated using the Ingenuity Pathway Analysis Molecule Activity Predictor (IPA MAP, Qiagen; Content Version 165342674) to predict the downstream functional consequences of modulating their expression.

#### Drug perturbation enrichment analysis

Drug perturbation enrichment analysis was performed using Enrichr ^33,34^ with the *LINCS L1000 Chemical Perturbations Up*^49^ library to identify compounds whose transcriptional response signatures significantly overlapped the spared neuron-specific transcriptional program. The 705 genes upregulated in the spared-specific transcriptional signature were used as input for the analysis. Enrichment significance was assessed using Enrichr combined scores and Benjamini–Hochberg-adjusted P values. Candidate compounds were ranked according to enrichment score, and the highest-ranked drug was selected for experimental validation.

#### Generation of hiPSC-derived glutamatergic neurons

Previously derived and validated passage-matched cryobanked induced neural progenitor cells (iNPCs) rapidly generated by transient overexpression of *NGN2*^87^ were used as a renewal source of hiPSC-derived cortical neurons to minimize inter-experimental variability. Briefly, control hiPSCs^77^ were transduced with lentiviral *pUBIQ-rtTA* and *tetO-NGN2-eGFP-NeoR*, subjected to 48-hour antibiotic selection with Geneticin (1mg/mL), and clonalized by expansion from single colonies. iNPCs were induced within 48-hours in “iNPC Induction Media”: DMEM/F12 with Glutamax, Glucose (0.3% v/v), N2 Supplement (1:100), Doxycycline (2 μg/mL), Geneticin (0.5 mg/mL), LDN-193189 (100 nM), SB431542 (5 μM), and XAV939 (1 μM), supplemented with Chroman I (25 ng/mL) for the first 24 hours. iNPCs are subsequently fed every 72-hours with “Expansion Media”: DMEM/F12 with Glutamax, Anti-Anti (1:100), MEM-NEAA (1:100), B27 minus Vitamin A (1:50), N2 Supplement (1:100), recombinant human basic FGF2 (20 ng/mL). iNPCs were split weekly by Accutase treatment for 10 minutes at 37°C and reseeded at a density of 10 x10^6^ cells onto Geltrex-(1:250) coated 10 cm dishes in DMEM.

To induce neurons from iNPCs, *NGN2* expression was re-induced with Doxycycline (1 μg/mL) at DIV 0 and selected with Geneticin (1mg/mL) for 72 hrs (DIV 1-4) and with FUdR (10 μM) for 48 hrs (DIV 2-4) in “Neuronal Induction Media”: DMEM/F12 with Glutamax, N2 Supplement (1:100), B27 minus Vitamin A (1:50). At DIV 4 iNPCs were dissociated with Accutase, seeded at a density of 1.0 x10^5^ cells per well onto 96-well plates coated with Poly-D-Lysine-(1:1) plus Natural Mouse Laminin-(1:100) in UltraPure water. Critically, to resolve differences in axon regeneration across genetic and pharmacological conditions, neurons were cultured in minimal maintenance medium lacking standard neuronal growth factors. “Neuronal Minimal Media”: Brainphys, Glutamax (1:100), Sodium Pyruvate (1 mM), Anti-Anti (1:100), N2 Supplement (1:100), B27 minus Vitamin A (1:50). Half media changes were performed every 72 hrs with Doxycycline (1 μg/mL) added until DIV 35 to maintain eGFP for live imaging^90^.

#### Neurite regeneration assay in hiPSC-neurons

Assay was adapted from our high-content imaging assay for neurite outgrowth^91^. Here, mechanical transection was performed at DIV 35 using the Incucyte® WoundMaker Tool (Sartorius, 4563). For genetic perturbations, neurons were pre-transduced with lentiviral shRNA constructs targeting *ARHGEF12* or a non-targeting control at DIV 30, at a multiplicity of infection (MOI) of 500, based on number of neurons detected per well by high-content imaging at the time of transduction. Lentiviral particles were generated in-house from MISSION shRNA plasmids (MilliporeSigma) as previously described ^92^. For *ARHGEF12*, 3 shRNA plasmids were pooled prior to lentiviral production, with target sequences CCACTCAAATGCAAAGGCTTA, GCGAGTATCCAGAGAAGGAAT, and GCGTTGCGTAATCATCCAGAA. The non-targeting control lentivirus was generated from a single shRNA plasmid with the target sequence GCGCGATAGCGCTAATAATTT. For drug perturbation experiments, neurons were treated immediately following injury at DIV 35 with either vehicle control or 1 μM vorinostat in Neuronal Minimal Media with Geltrex (1:62.5).

Neurite regeneration was quantified by daily high-content imaging for 3 days using a Thermo Scientific CellInsight CX7 LZR High-Content Screening Platform with a 10X objective. Neurons were visualized in the 488-nm channel by fluorescence from the NGN2-eGFP construct. Automated neuron quantification and neurite tracing was performed from the scraped area using the Neuronal Profiling v4.2 application in Thermo Scientific HCS Studio Cellomics Scan v6.6.2. For drug perturbation experiments, 15 wells were imaged per condition, whereas 30 wells per condition were imaged for genetic manipulation experiments. Nine tiled images were acquired per well to comprehensively examine the entire surface; each “n” therefore represents the total number of neurons and total neurite length per well, derived from thousands of neurons. Prior to quantification, images underwent quality control and wells with substantial variation in scrape width, extensive cell detachment, or evidence of unhealthy cultures were excluded. Total neurite length was normalized to baseline neuron count using a condition-specific linear mixed-effects model and analyzed using linear mixed-effects models as described in the Quantification and statistical analysis section. Total neurite length was normalized using a linear mixed-effects model incorporating baseline neuron count as a covariate. Normalized neurite length was subsequently analyzed using a linear mixed-effects models accounting for condition, imaging day, their interaction, and repeated measurements within wells.

#### Gene expression analysis by qPCR

All molecular biology and quantitative PCR procedures were performed according to the manufacturers’ protocols. On day 38, neurons were harvested, wells from each condition were pooled into five independent biological samples, and total RNA was isolated using the PicoPure RNA Isolation Kit. First-strand cDNA was synthesized using the SuperScript III First-Strand Synthesis System. Quantitative PCR was performed using Power SYBR™ Green PCR Master Mix with 100 ng cDNA and 1 μM primers. Relative gene expression was quantified using the 2^−ΔΔCt method with GAPDH as the reference gene. n = 5 per condition. Primer sequences were as follows: *ARHGEF12* forward, AGGAACAGACGCTGGATACCTG; reverse, GTGTGCCATCTAAGGTGTCTCC; GAPDH forward, GTCTCCTCTGACTTCAACAGCG; reverse, ACCACCCTGTTGCTGTAGCCAA.

### Quantification and statistical analysis

Statistical analyses were performed in R (v4.3.3) and Python (v3.12.9). Differential expression analysis was performed using PyDESeq2 as described above. Genes with a padj ≤ 0.05 and |Log2(FC)| ≥ 0.25 were considered significantly differentially expressed and used for all downstream functional analyses. For Ingenuity Pathway Analysis (IPA), canonical pathways with Benjamini–Hochberg false discovery rate (FDR) ≤ 0.05 were considered significantly enriched and used for downstream analyses and visualization. Diseases and biological functions and upstream regulators with enrichment p ≤ 0.05 were considered significant. For Enrichr transcription factor enrichment analysis, transcription factor terms with a padj ≤ 0.05 were considered significantly enriched.

Neurite regeneration data were analyzed in R (v4.3.3) using linear mixed-effects models implemented with the lme4 (v1.1.37) and emmeans (v2.0.3) packages. Experimental condition, imaging day, and their interaction were included as fixed effects, whereas well was modeled with random intercepts to account for repeated measurements. Pairwise comparisons between experimental conditions and the control were performed at each imaging day using estimated marginal means. Unless otherwise specified, all statistical tests were two-sided, data are presented as mean ± SEM, and p-values were adjusted for multiple comparisons using FDR correction, as appropriate. Exact sample sizes, statistical tests, and p-values and adjusted p-values are reported in the corresponding figure legends.

